# Morphological and physiological response strategies of *Vallisneria natans* at different water depths and light conditions

**DOI:** 10.1101/2020.03.20.999904

**Authors:** Qingchuan Chou, Jianfeng Chen, Wei Zhang, Wenjing Ren, Changbo Yuan, Xiaolin Zhang, Te Cao, Leyi Ni, Erik Jeppesen

**Author notes:** **Author email addresses:** Qingchuan Chou, Jianfeng Chen, Wei Zhang, Wenjing Ren, Changbo Yuan, Xiaolin Zhang, Te Cao, Leyi Ni, Erik Jeppesen. **Corresponding author:** Xiaolin Zhang, Donghu Experimental Station of Lake Ecosystem, State Key Laboratory of Freshwater Ecology and Biotechnology, Institute of Hydrobiology, The Chinese Academy of Sciences, Wuhan 430072, China.

## Abstract

Phenotypic plasticity is an important adaptation to spatial and temporal environmental variations. For submerged macrophytes, adaptation to water depth and light variation is particularly important. To determine the morphological and physiological adaptive strategies of *Vallisneria natans* at different water depths and light conditions, we combined field investigation, light control experiment and *in situ* physiological response experiment. In the field investigation and the light control experiment, both water depth and light intensity had prominent effects on the morphological of *V. natans*, especially in fresh weight and leaf length. The leaf length elongated more rapidly at intermediate water depth sites with lower light intensity. In the *in situ* experiment, the survival boundary of *V. natans* is 5.5 m in Lake Erhai. Below this depth, the chlorophyll-a content increased gradually with increasing water depth. Our results demonstrated that *V. natans* can adapt to water depth and light availability by changing morphological, physiological and resource allocation. At low light condition, *V. natans* invested more resource for light acquisition, simultaneously, changing the photosynthetic pigment content to compensate for light attenuation; conversely, more resource was directed towards reproduction. These results will provide new insight for species selection when conducting aquatic plants restoration in freshwater ecosystem.

**HIGHLIGHTS:**

- Water depth and light availability affect the morphology, physiology, and resource allocation of *V. natans*.
- An alternative resource allocation pattern of *V. natans* could shift between light acquisition and reproduction.

## Introduction

Phenotypic plasticity is the ability of a genotype to modify its phenotype in response to environmental changes as a consequence of an interaction between genes and the environment (Bradshaw, 1965; 2006). Plasticity is the potential of organisms to produce a range of different and suitable phenotypes in a complex environment (Dewitt et al., 1998). Phenotypic plasticity is ubiquitous in all types of organisms (Sultan, 1987; Sultan, 1995; Tollrian, 2002) and is an important ecological strategy for plants adapt to heterogeneous habitats (Sultan, 1995; Orr, 1999; Huey et al., 2000; Debat and David, 2001; Mai and Lovett-Doust, 2005; Bradshaw, 2006). As emphasized in life history theory, shifts in phenotypic plasticity can be instrumental for maintaining plant fitness in a changing environment (Li et al., 2018). Phenotypic plasticity increases resistance and adaptability and allows species to have a wider ecological niche, to better tolerate stressful environmental conditions, and to expand their potential range of available diet resources (Sultan and Bazzaz, 1993; Cao et al., 2016). Thus, phenotypic plasticity allows species to occupy a wider geographical range and more diverse habitats (Burns and Winn, 2006), which can buffer against and to some extent reduce the selective pressure caused by the new habitat, and according to the niche theory a specialist may evolve into a generalist (Sultan and Bazzaz, 1993; Sultan, 1995; Sultan, 2000). Natural habitat heterogeneity makes it difficult for plants to acquire resources (Jackson and Caldwell, 1993; Chen et al., 2002), and many plants therefore exhibit high phenotypic plasticity, allowing them to adapt to heterogeneous habitats. Plants with high plasticity therefore have a distinct competitive advantage over species with low plasticity (Sultan, 1995; Orr, 1999; Bradshaw, 2006). Phenotypic plasticity may affect the different life stages of the plants via various mechanisms through which morphological and physiological countermeasures to environmental stress are developed and manifested in a wide diversity of patterns (Brewer, 1999; Via and Hawthorne, 2002).

As one of the most important primary producers, submerged macrophytes are important for the ecological state and diversity in many aquatic ecosystems. Many submerged macrophytes have low genetic diversity but occupy diverse habitats. Phenotypic plasticity therefore plays an important role in the adaptation of the submerged macrophytes to heterogeneous habitats (Bradshaw, 1965). Submerged macrophytes generally adapt to heterogeneous habitats by plasticity strategies (morphological and physiological plasticity) (Richards, 1968; Grime et al., 1986; Sultan, 2000; Garbey et al., 2006) to maximize acquiring resources for growth and reproduction (Ikegami et al., 2008; Wang et al., 2011). Such strategies can greatly reduce environmental stress and improve their adaptability to heterogeneous environments (Schlichting, 1986; Tollrian, 2002).

Water depth and underwater light intensity are crucial to the survival and growth of submerged macrophytes. Light availability is severely attenuated by water depth, which may be the main factor restricting the growth of submerged macrophytes as photosynthesis can be constrained in deep water (Li et al., 2018). Along with the increasing water depth, the underwater light intensity decreases dramatically, submerged macrophytes usually show adaptations through a variety of adjustments of plant morphology, biomass reallocation, photosynthesis pigment content, and root traits (Maberly, 1993; Strand, 2001; Hussner et al., 2009). Many studies have investigated the phenotypic plasticity of submerged macrophyte species that exhibit high phenotypic plasticity in response to changes in water depth and light intensity (Yang et al., 2004; Geng et al., 2007; Hyldgaard and Brix, 2012; Riis et al., 2012; Zhu et al., 2012; Malheiro et al., 2013; Reckendorfer *et al*., 2013; Atapaththu and Asaeda, 2015; Baastrup-Spohr et al., 2016; Wei et al., 2018). The availability of light under water is an important factor determining the maximum distribution depth of macrophytes in a proper aquatic ecosystem (Zhang et al., 2007; Liu et al., 2016). One of the important species for ecological restoration of the water environment in subtropical and tropical lakes (Ke and Li, 2006; Qin et al., 2013) is *Vallisneria natans* whose life history is also affected by water depth and light intensity. In this study, our principal aims are to determine the emergency remediation mechanism of *V. natans* at different water depths and light conditions, including morphological and physiological responses, based on field investigations, a light control experiment and an *in situ* physiological responses experiment. We hypothesized that: (1) *V. natans* would allocate more biomass for rapid elongation of leaves to obtain an optimal size for light harvesting and thus alleviate low light stress in deeper and low light intensity habitats; (2) as an auxiliary measure to morphological plasticity, *V. natans* would also develop special physiological adaptive mechanisms changing the photosynthetic pigment content to compensate for light attenuation in deeper water.

## Materials and Methods

### Species description

*Vallisneria natans* L. (Hydrocharitaceae) is a perennial submerged clonal freshwater macrophyte. It has many vertical fibrous shoots from the base of its short stem (Lowden, 1982; Mcconchie and Kadereit, 1987). Its leaves are submerged or floating and linear shaped (Lowden 1982), and it has an elongate axis with both stolon and leaf-bearing branches (Mcconchie and Kadereit, 1987).

*V. natans* commonly inhabits lakes, swamps, ponds, rivers, ditches, and other water areas. It adapts well to water depth, water quality, sediment, etc. Its morphological characteristics often change with the water depth and it may reach high abundances in favorable environments (Chen et al., 2012).

*V. natans* is widely distributed in tropical and subtropical climates, including North America, Europe, Africa, Asia, Oceania, and Australia (Lowden, 1982; Mcconchie and Kadereit, 1987).

### Field investigation

The field investigation was conducted from 1-23 September 2016 in Lake Erhai, Yunnan Province, China. We collected submerged macrophytes using a reaping hook (covering 0.2 m^2^) along a gradient of five water depths (WD): 0.5 m, 1.5 m, 2.5 m, 3.5 m, and 4.5 m. We collected a total of 786 submerged macrophyte samples (n) from 16 sampling sites (Fig. 1) located around the lake shore. The plants were washed carefully to remove mud, attached algae, eggs of fishes, and zoobenthos. We then measured: (1) fresh weight of the plant (FW), (2) length of the longest leaf (LL), (3) length of the longest root (RL), and (4) leaf number (LN).

**Fig. 1.**
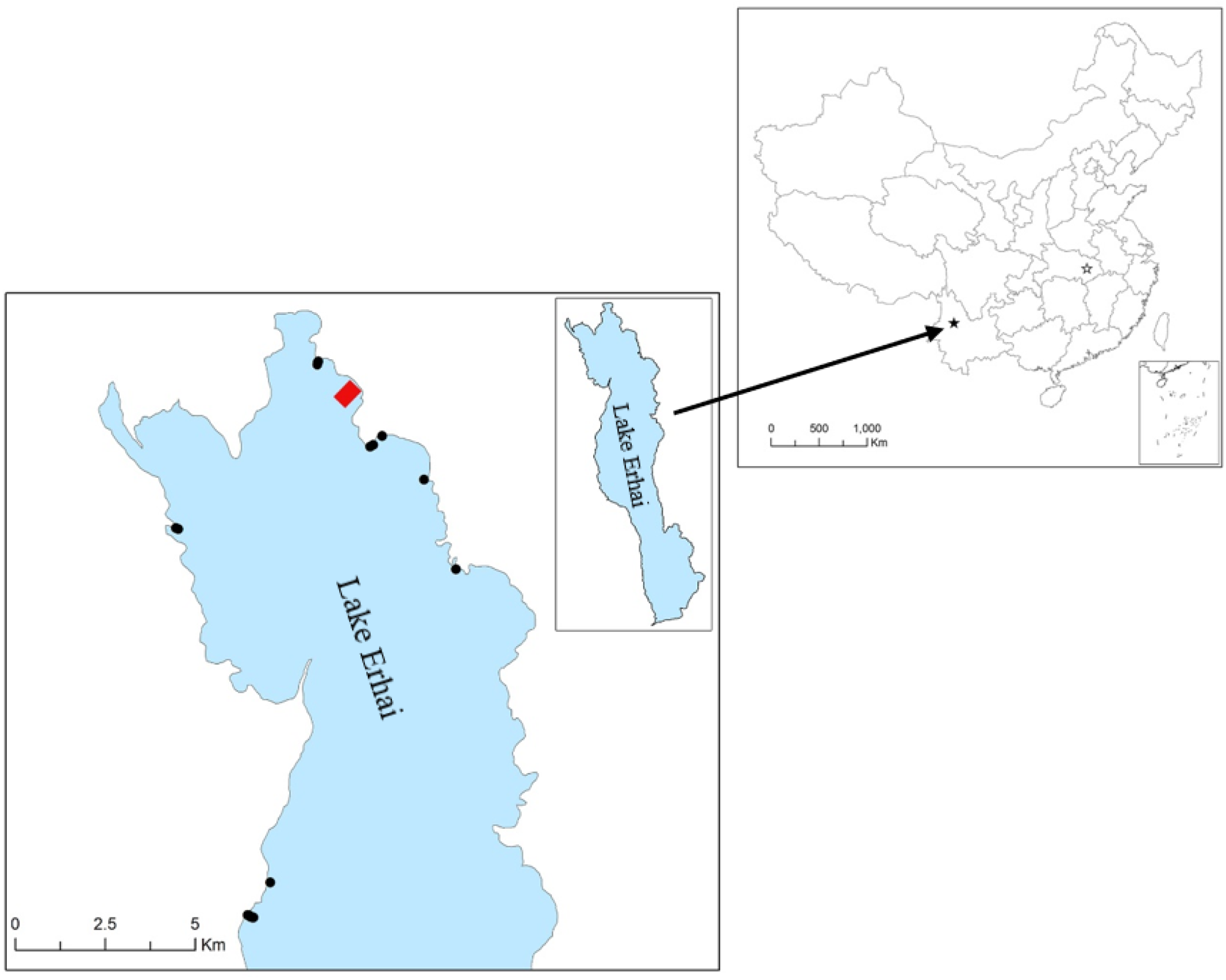
Map showing field investigation sampling sites and location of light control and physiological response experiments. Black solid pentagram = Lake Erhai; black hollow pentagram = light control experiment, Donghu Experimental Station, Wuhan, China; black solid dots = field investigation sampling sites; red area = location of in situ physiologic response experiment.

During the investigation, major physical and chemical parameters of the water column were measured (temperature (Temp), total nitrogen (TN), total phosphorus (TP), and chlorophyll a (Chl-a) based on the methods described by APHA (2012). Surface sediments were collected from the top 10 cm layer with a Peterson sediment collector, the sediment nitrogen (SN) concentrations were determined by an elemental analyzer (Flash EA 1112 series, CE Instruments, Italy) and the sediment phosphorus (SP) concentrations were determined following the method described by Bao (2000). Photosynthetically active radiation (PAR) was measured at noon on sunny days by a Li-COR sensor coupled with a data logger (Li-1400; Li-Cor Company, Lincoln, NE, U.S.A.). Detailed information about Temp, TN, TP, Chl-a, SN, and SP at the sampling sites is given in Table S1.

### Light control experiment

The light control experiment was performed at the Donghu Experimental Station of Lake Ecosystem, Institute of Hydrobiology, Chinese Academy of Sciences, from 20 April 2015 to 20 April 2016. The seedlings of *V. natans* (31.7±8.7 cm height, mean ± SD) and the sediment for the experiment were collected from Lake Donghu. Seedlings with almost the same height were chosen and placed in an experimental material pool to ensure the same initial status of the experimental material. Three randomly selected seedlings from the material pool were planted in each outdoor glass aquarium (length: 50 cm; width: 50 cm; height: 80 cm), which contained 10 cm sediment. The water used in the experiment was a mixture of lake water and tap water (with a volume ratio as 3:7). The experiment designed four different light treatments: (1) 39.5% (L1), (2) 17.1% (L2), (3) 7.1% (L3) and (4) 2.8% (L4) of full daylight. Sheltering was obtained using a one-layer black nylon mesh (L1), a two-layer black nylon mesh (L2), a three-layer black nylon mesh (L3), and a four-layer black nylon mesh (L4), which did not alter the different wave-length ratios of the incident light. Each group had three replicates. The plants were harvested after 12 months and weighed to determine fresh weight (FW), leaf length (LL), and root length (RL), and the number of leaves (LN) was counted.

During the light control experiment, PAR, Temp, TN, TP, and Chl-a in the water column were monitored every two weeks based on the same methods and equipment as field investigation. More detailed information about PAR, Temp, TN, TP, and Chl-a concentrations in the light control experiment is given in Table S2.

The light extinction coefficient (K) of Lake Erhai was calculated based on the equation: K = (lnI1-lnI2)/(d2-d1), where d_1_ and d_2_ stand for water depth, and I_1_ and I_2_ are PAR at water depth d_1_ and d_2_, respectively (Chen et al., 2016). Then, we calculated the corresponding water depth according to the specific sheltering light intensity and the K value of Lake Erhai. The calculated water depths for the different light treatments were 0.66 m, 1.52 m, 2.52 m, and 3.5 m for L1, L2, L3, and L4, respectively.

### *In situ* physiological response experiment

The *in situ* physiological response experiment was conducted at the Aoshan Bay (located in the north of Lake Erhai), from 18 September to 18 November in 2016. A total 450 *V. natans* seedlings with the same height and biomass (13.5±1.6 cm height, 1.1±0.3 g fresh weight, mean ± SD) were collected from the submerged macrophyte nursery at the Erhai Lake Research Station for this experiment. Mud from the 10 cm part of the sediment collected in Aoshan Bay was used. The experiment was divided into nine different water depth gradients (0.5 m, 1.5 m, 2.5 m, 3.5 m, 4.5 m, 5.5 m, 6.5 m, 7.5 m, and 8.5 m). Every 10 seedlings were planted in a non-transparent plastic bucket (inner diameter: 35 cm, inner height: 26 cm) filled with 20 cm mud. The plastic buckets were then suspended in the experimental area (Fig. 1, with a maximum water depth above 9 m). Five replicates were made for each water depth. The plants were harvested after 60 days and carefully washed to remove mud and attached algae, after which they were weighed to determine FW and analyzed for chlorophyll a (Chl-a) content with a spectrophotometer following liquid nitrogen extraction (Murata et al., 1973).

During the *in situ* physiological response experiment period, PAR, Temp, TN, TP, and Chl-a in the water column of the experimental area were measured every two weeks applying the same methods and equipment as field investigation. More detailed information about PAR, Temp, TN, TP, and Chl-a concentrations in the *in situ* physiological response experiment can be found in Table S3.

### Statistical analyses

We used IBM SPSS 22.0 for one-way analysis of variance (ANOVA) to test the differentiation in water quality during the experimental period. The differences in environmental factors and morphological parameters between the sampling sites in the field investigation and the different treatments in the light control experiment were analyzed using principal component analysis (PCA) within Canoco 5.0 (Fig. 2 and 3). In the field investigation and light control experiment, the effects of water depth and light intensity gradients on FW, LL, RL, and LN of *V. natans* L. were analyzed using one-way ANOVA (Table S4 and S5), and the relationship between FW, LL, RL, and LN at different water depths and light intensities was analyzed with linear regression in OriginPro 9.0 (Fig. 4, 5 and 6). We used the psych package of R to run Spearman correlation analyses of WD, PAR, FW, LL, RL, and LN in the field investigation and the light control experiment (Fig. 7). Finally, we used OriginPro 9.0 to analyze changes in FW, the relative growth rate (RGR), and the leaf chlorophyll-a content of *V. natans* L. at different water depths in the *in situ* physiological response experiment (Fig. 8).

**Fig. 2.**
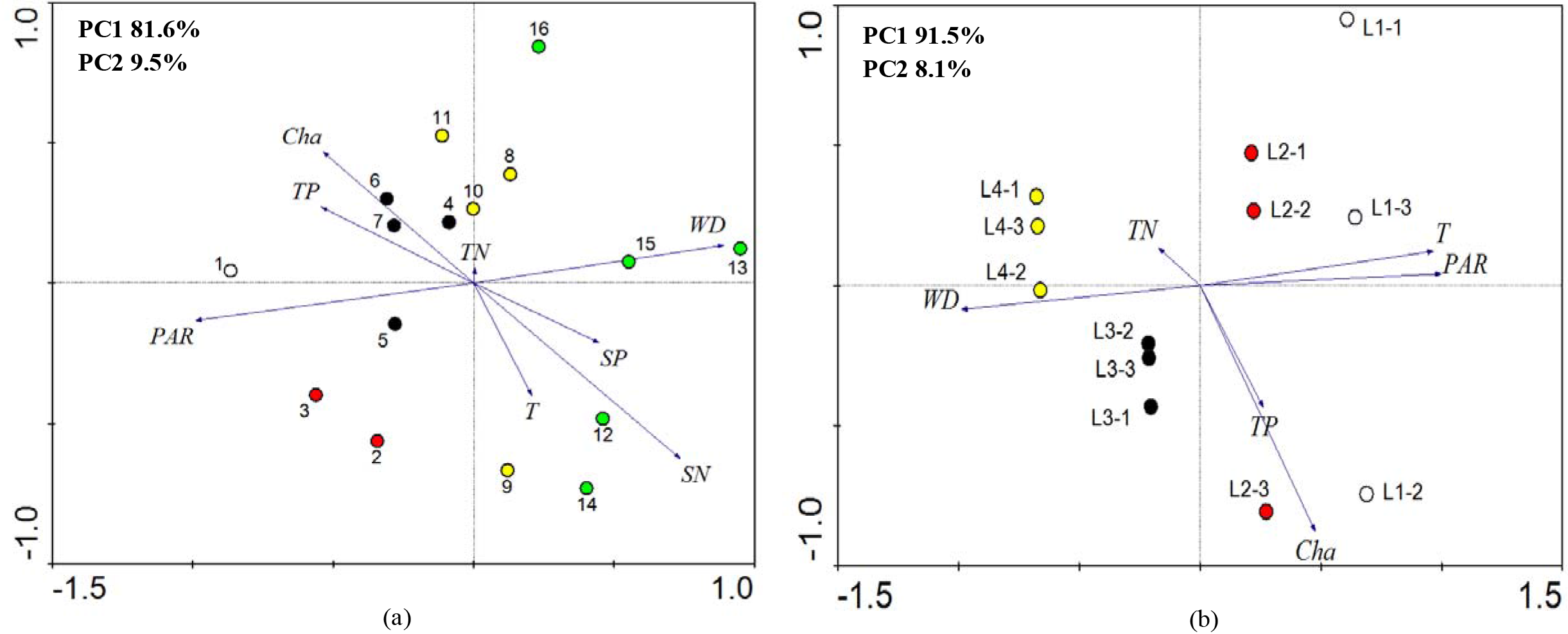
Environmental indicators in the PCA analysis of the results of the field investigation (a) and the light control experiment (b). Same colors in the field investigation and the light control experiment mean that water depth and light treatment gradients were identical. WD = water depth; PAR = photosynthetically active radiation; T = temperature; TN = total nitrogen in water; TP = total phosphorus in water; Chl-a = chlorophyll a in water; SN = total nitrogen in sediment; SP = total phosphorus in sediment.

**Fig. 3.**
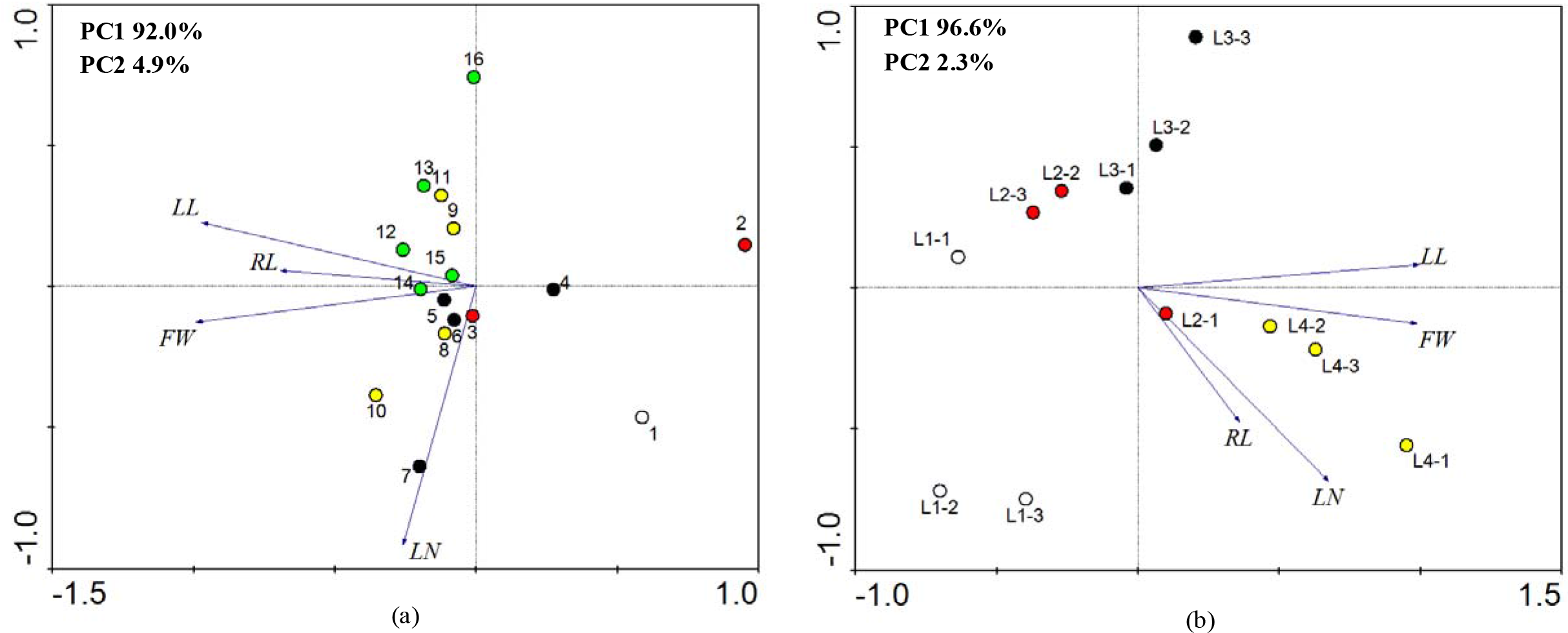
Morphological indicators in the PCA analysis conducted for the field investigation (a) and the light control experiments (b). Same colors in the field investigation and the light control experiment mean that water depth and light treatment gradients were identical. FW = fresh weight; LL = leaf length; RL = root length; LN = leaf number.

**Fig. 4.**
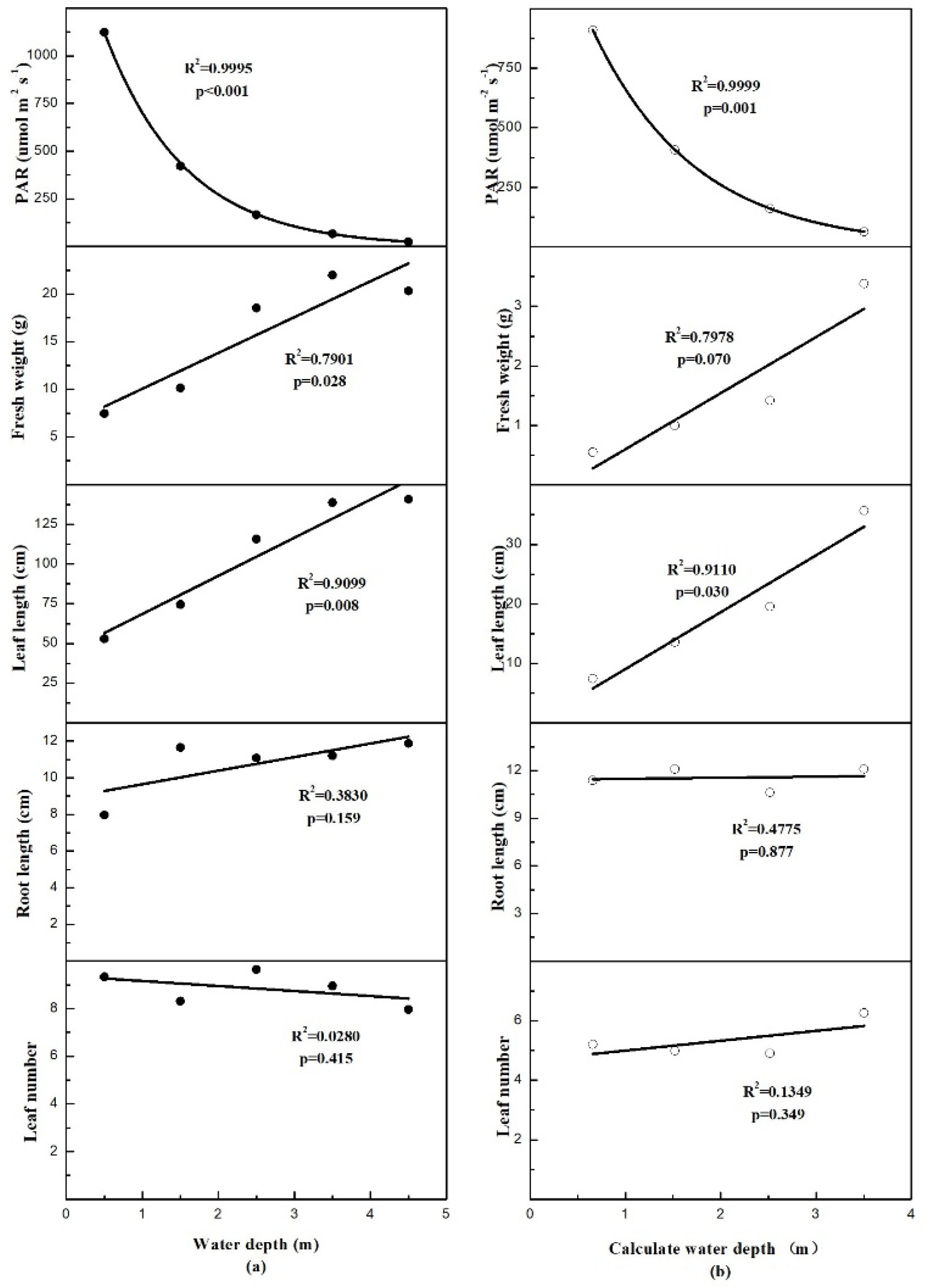
The relationships between FW, LL, RL and LN of V. natans L. along a water depth gradient in the field investigation (a) and the light control experiment (b).

**Fig. 5.**
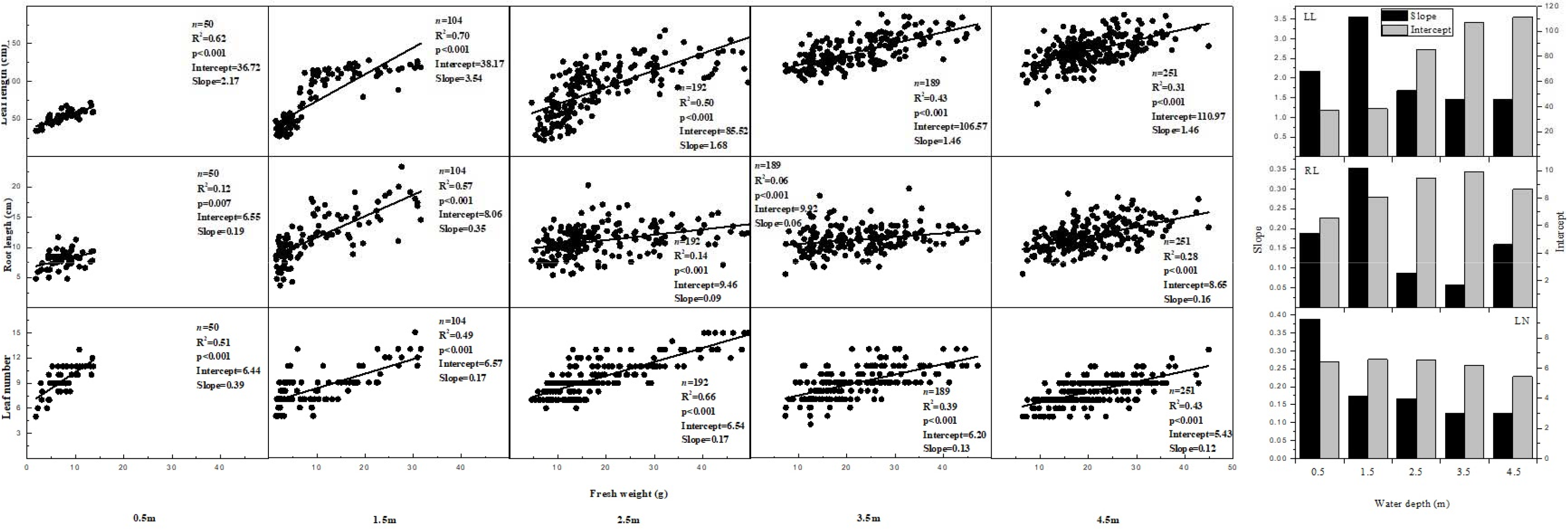
Relationship between fresh weight (FW) and leaf length (LL), root length (RL), and leaf number (LN) of *Vallisneria natans L*. at different water depth gradients and variation of slope and intercept of morphological parameters of *V. natans L*. with water depth gradient in the field investigation.

**Fig. 6.**
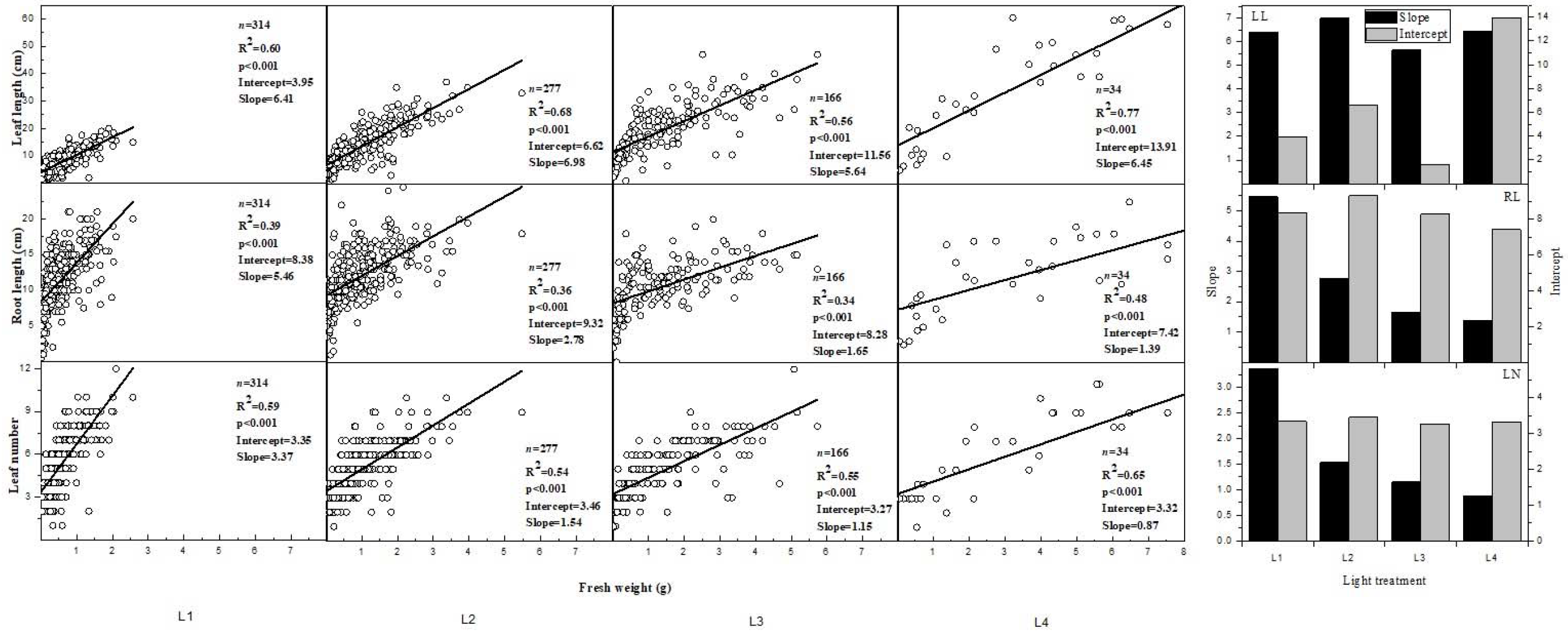
Relationship between fresh weight (FW) and leaf length (LL), root length (RL), and leaf number (LN) of *Vallisneria natans L*. at different light intensity gradients and variation of slope and intercept of morphological parameters of *V. natans L*. with light intensity gradients in the light control experiment.

**Fig. 7.**
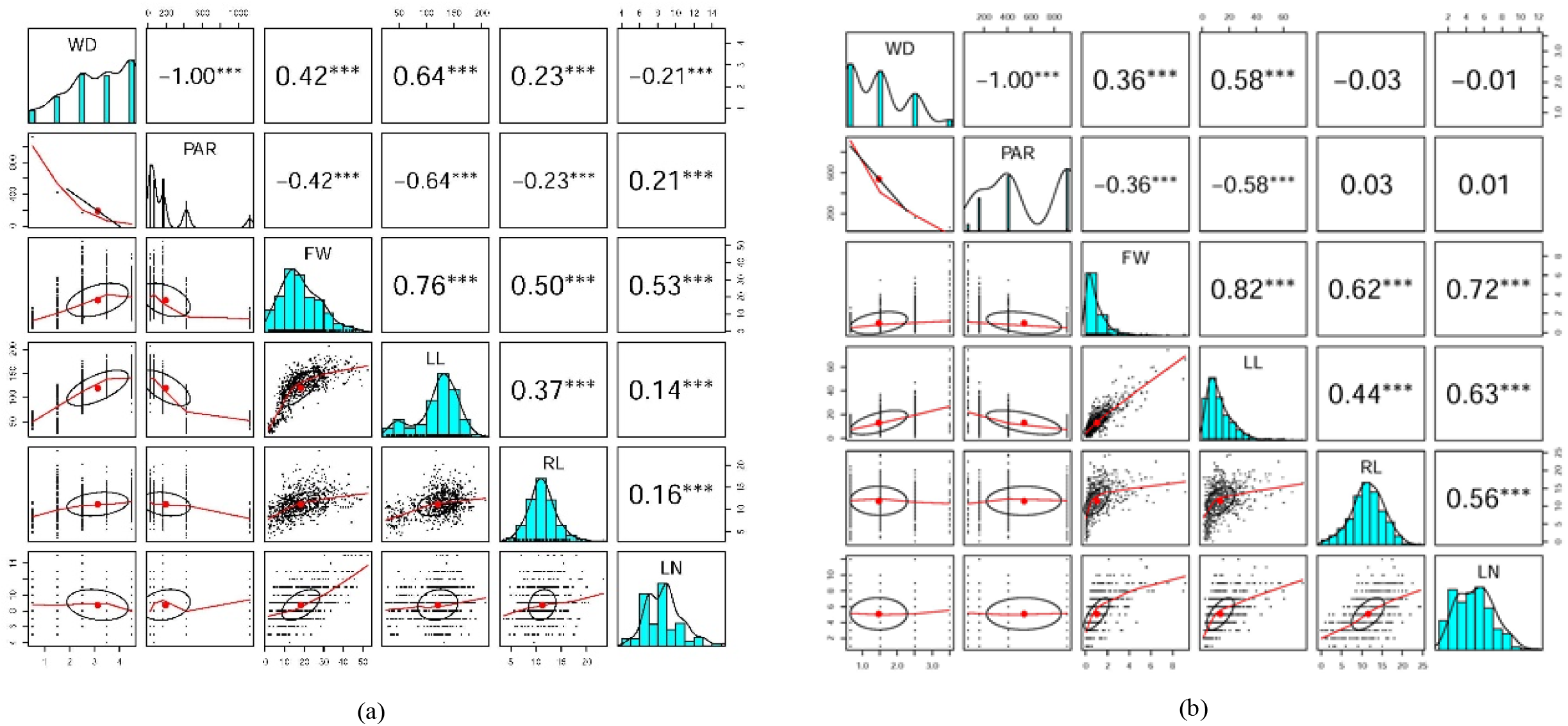
Spearman correlation analysis between WD, PAR, FW, LL, RL, and LN in the field investigation (a) and light control experiments (b). * P<0.05, ** P<0.01, *** P<0.001, and without asterisk P>0.05. WD = water depth; PAR = photosynthetically active radiation; FW = fresh weight; LL = leaf length; RL = root length; LN = leaf number.

**Fig. 8.**
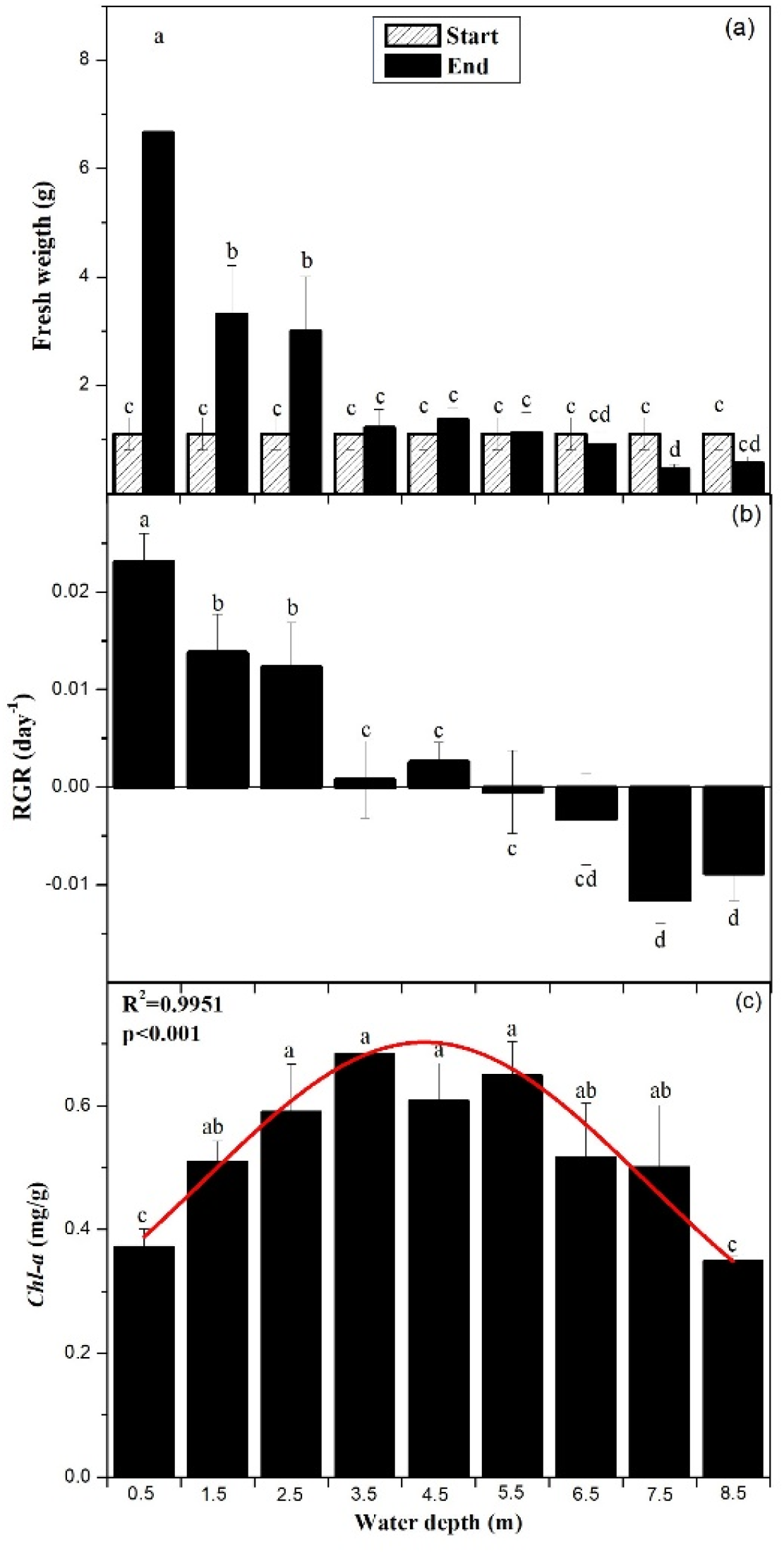
Average ramet biomass at the start and at the end of the experiment, average relative growth rate (RGR), and changes in chlorophyll-a levels along a water depth gradient in the *in situ* physiological response experiment.

## Results

### The main environmental factors affecting the morphological plasticity of *V. natans*

One-way ANOVA showed that there were no significant differences (P>0.05) in Temp, TN, TP, and Chl-a but a significant difference (P<0.05) in PAR results between the field investigation and the light control experiment (Table S1 and S2). Based on the PCA results (Fig. 2 a, b), nonlinear fitting of PAR and water depth (Fig. 4 a, b), and Spearman correlation analysis (Fig. 7 a, b) of the field data and data from the light control experiment, we found that water depth (WD) and PAR were the major environmental factors affecting the morphological plasticity of *V. natans*. In the field investigation, the PCA analysis showed that different sampling sites can be distinctly separated by WD and PAR, the eigenvalues of Axis 1 and Axis 2 were 81.6% and 9.5% (Fig. 2 a), respectively, indicating that 91.1% of the total variance could be explained by the two axes. In the light control experiment, the PCA also identified WD and PAR as most important, the eigenvalues of Axis 1 and Axis 2 being 91.5% and 8.1% (Fig. 2 b), respectively. Thus, 99.6% of the total variance could be explained by the two axes. Nonlinear fitting indicated that PAR, as expected, decreased exponentially with increasing water depth (Fig. 4 a, b).

### Morphological response of *V. natans* along water depth and light gradients

Water depth and light intensity also had prominent effects on plant size. In the field investigation and the light control experiment, the morphological indicators FW and LL showed the most significant changes with water depth and light intensity (Fig. 3 a, b, and Fig. 4 a, b). Four morphological parameters (FW, LL, RL, and LN) were analyzed (Table S4 and S5). In the field investigation, all morphological indicators correlated significantly (p<0.001) with water depth (Fig. 7 a). In the light control experiment, FW and LL were significantly correlated (p<0.001) with light intensity, while RL and LN did not exhibit any significant correlation (p>0.05) with these two variables (Fig. 7 b). Using linear regression, we found that the fresh weight and leaf length of each ramet were significantly (p<0.05) higher and longer in the deeper water and at low light intensity, whereas root length and leaf number demonstrated no significant variation (p>0.05) (Fig. 4 a, b).

### Correlation of various morphological indicators

Significant positive correlations (p<0.001) were found between all morphological indicators studied in both the field investigation and the light control experiment (Fig. 7 a, b). As fresh weight is the concentrated expression of leaf length, root length, and leaf number, we used fresh weight as the abscissa and leaf length, root length, and leaf number as ordinates to analyze the trends of changes with alterations in fresh weight at different gradients of water depth and light intensity.

The LL-FW, RL-FW, and LN-FW allometric relationships were significantly related to water depth and light intensity but had different slopes (Fig. 5 and 6). In the field investigation and light control experiment, the LL-FW slopes first increased and then gradually decreased when the water depth increased and the light intensity decreased, while the RL-FW and LN-FW slopes decreased, indicating a biomass allocation favoring leaf growth rather than root and leaf number growth. In the field investigation, leaf length and fresh weight were closely correlated; thus, with increasing fresh weight, the leaf length increased as well, and the allometric growth rate increased with an increasing water depth. Root length also varied relative to fresh weight, root length increasing with the increasing fresh weight; however, the allometric growth rate decreased with increasing water depth. Also, leaf number was affected by the fresh weight and increased with increasing fresh weight, and the allometric growth rate declined with rising water depth (Fig. 5). In the light control experiment, leaf length exhibited a significant positive correlation with fresh weight and the allometric growth rate increased with the continuous weakening of the light intensity. Root length also demonstrated a significant positive correlation with fresh weight, the allometric growth rate declining with diminishing light intensity. Fresh weight also had a significant positive relation with leaf number whose allometric growth rate decreased with decreasing light intensity (Fig. 6). Therefore, *V. natans* at different water depths and light intensity differed in their allometric relationships.

### Physiological responses of *V. natans* along a water depth gradient

In the *in situ* physiological response experiment, one-way ANOVA showed that there were no significant differences (P>0.05) in Temp, TN, TP, and Chl-a but a significant difference (P<0.05) in PAR results (Table S3), while increasing water depth influenced the relative growth rate (RGR) and photosynthetic pigments content of *V. natans* (Fig. 8). At the end of the *in situ* physiological response experiment, the mean fresh weight of the plants in the 0.5 m, 1.5 m, and 2.5 m were significantly (p<0.05) higher than the initial weights, while there was no significant (p>0.05) difference in mean fresh weight between the 3.5 m, 4.5 m, and 5.5 m ramets. In the 6.5 m, 7.5 m, and 8.5 m ramets, fresh weight was significantly (p<0.05) lower than the initial weights (Fig. 8 a). When water depth was greater than 5.5 m, RGR became negative, implying that the experimental *V. natans* seedlings did not survive for long at water depths > 5.5 m. In Lake Erhai, the water depth boundary for *V. natans* survival is 5.5 m (Fig. 8 b), as same as the light compensation points depth. The photosynthetic pigment (chlorophyll-a) content of *V. natans* first typically increased and then gradually decreased with the increase in water depth, non-linear fitting showing that the turning point at which the concentration of photosynthetic pigments began to decrease was 5.5 m where RGR became negative (Fig. 8 c).

### Population expansion strategy affected by light intensity

The vegetative growth strategy of V. *natans* at various water depths and light conditions differed. The population growth of *V. natans* was high at high light conditions and vice versa. In the light control experiment, light intensity exhibited significant effects on the reproductive strategy of *V. natans*, the number of ramets counted at the end of the light control experiment were (means ± SD) 105 ±40^a^, 92 ± 3^b^, 55 ± 14^c^, and 11 ± 2^d^ in L1, L2, L3, and L4, respectively (Table S5). Most ramets appeared in the L1 treatment where the leaves of *V. natans* grew slowly and more energy was allocated to expansion rather than to leaf growth. As the light intensity decreased, *V. natans* allocated more energy to leaf growth, and the lowest number of ramets appeared in the L4 light treatment.

## Discussion

Our study clearly showed that phenotypic plasticity is an important mechanism/strategy for *V. natans* to adapt to the changes of water depth and light intensity. Aquatic plants typically adjust their organ biomass allocation and morphology to improve their ability to exploit critical resources (Westoby et al., 2002; Xiao et al., 2007; Xie et al., 2007; Poorter et al., 2012). In our study, *V. natans* had the lowest fresh weight and shortest leaf length in the shallow waters and in the high light treatment. The fresh weight and leaf length increased progressively with increasing water depth and decreasing light intensity. These results are consistent with our first hypothesis. Shallow water and high light intensity may inhibit the development of *V. natans* and reduce plant growth (Simpson et al., 1980; Bonser and Geber 2005; Weijschedé et al., 2006; Lu et al., 2013; Cao et al., 2016). This inhibitory effect gradually disappeared as the water depth increased and light intensity decreased, leading to higher plant height and allocation of more biomass to elongation of leaves while reduce root growth. In this way, leaves may more rapidly reach the upper layer to achieve better light conditions. This light harvesting strategy, called ‘light foraging’, allows plants to obtain light at minimum metabolic cost (Hutchings and Kroon 1994). Others have also found that *V. natans* adopts plastic strategies and increases its leaf biomass and leaf length to reach a given plant size in response to increasing water depth and decreasing light intensity at the expense of branch number and belowground biomass (Lieffers and Shay, 1981; Nohara and Kimura, 1997; Vretare et al., 2001). However, Fu (2012), inconsistently with our results, found small-sized *V. natans* individuals in deep and low light intensity waters. Their experiments lasted only 52 days, which perhaps was a too short period to allow the plants to fully adapt to the new habitat.

Besides morphological adaptation, physiological plasticity may also occur (Anthony and Fabricius, 2000; Hoogenboom et al., 2008). Changing the content of photosynthetic pigments is the main physiological adaptive mechanism (Cao et al., 2016). In agreement with our second hypothesis, we found that water depth had a significant effect on the photosynthetic pigment (chlorophyll-a) of *V. natans*. It increased with increasing depth until 5.5 m and then gradually decreased at deeper sites. *V. natans* may shift the light compensation point to cope with reduced light (Bai et al., 2015; Wei et al., 2018). Blanch (1998) found that *V. natans* had a low photosynthetic light intensity compensation point of 9.4 umol m^−2^ s^−1^, which is close to the light intensity value (9.0 umol m^−2^ s^−1^) at 5.5 m water depth in our *in situ* physiological response experiment. Accordingly, compared with the field survey data of light intensity at 4.5m, we concluded that the survival boundary for *V. natans* in Lake Erhai is 5.5m.

We found that *V. natans* faced a trade-off between biomass increase and clonal expansion. In the light control experiment, the population growth increased with light, which agrees with the trade-off theory (Thompson and Eckert, 2004; Reusch, 2006; Van Drunen and Dorken, 2012; Li et al., 2018). *V. natans* clonal growth and regeneration depend strongly on water depth and light availability (Ferreiro et al., 2013; Søndergaard et al., 2013; Dong et al., 2014; Fu et al., 2014; Wei et al., 2018). When light resources are sufficient, *V. natans* allocates more energy to reproduction and population growth as a more abundant populations can share or prevent the likelihood of extinction, while under low light conditions, *V. natans* allocated most of its biomass to growth of light resource acquisition organs and the least to reproduction (Li et al., 2018). Darkness therefore increases the likelihood of population extinction (Xiao et al., 2007).

In conclusion, our results evidenced that water depth and light availability are important driving factors for population maintenance and expansion of *V. natans*. And this submerged species evolved phenotypic plasticity to adjust morphological, physiological and also reproductive characters to better adapt different environments. At low light, the leaf length of *V. natans* grew faster and the root length and leaf number increased slowly as the plants used more energy to increase the organ for acquisition of light; in contrast, when light resources were sufficient, more energy was directed towards reproduction. Such phenotypic plasticity will shape the population characters of this species and further influence the local community assembly, this adaptive mechanism also provides new insight for species selection when conducting ecological restoration of lost aquatic plants in subtropical and tropical freshwater ecosystem.

## Acknowledgements

We are grateful to A.M. Poulsen for manuscript editing. This work was supported by the National Natural Science Foundation of China (31870446, 31200356), and the Special Foundation for State Basic Working Program of China (2013FY112300). It was also partly supported by the State Key Laboratory of Freshwater Ecology and Biotechnology (2019FBZ01) and a scholarship from China Scholarship Council (Grant No. 201704910729) to Q.C. Chou for a visit to Aarhus University. E. Jeppesen was supported by the AU Centre for Water Technology (watec.au.dk) and the Tübitak BIDEB 2232 program.

